# Recoding enhances the metabolic capabilities of two novel methylotrophic Asgardarchaeota lineages

**DOI:** 10.1101/2021.02.19.431964

**Authors:** Jiarui Sun, Paul N. Evans, Emma J. Gagen, Ben J. Woodcroft, Brian P. Hedlund, Tanja Woyke, Philip Hugenholtz, Christian Rinke

**Affiliations:** Australian Centre for Ecogenomics, School of Chemistry and Molecular Biosciences, The University of Queensland, St Lucia, Queensland, Australia; School of Earth and Environmental Sciences, The University of Queensland, St Lucia, Queensland, Australia; Centre for Microbiome Research, School of Biomedical Sciences, Queensland University of Technology (QUT), Translational Research Institute, Woolloongabba, Australia; School of Life Sciences and Nevada Institute of Personalized Medicine, University of Nevada, Las Vegas, Nevada, USA; DOE Joint Genome Institute, Walnut Creek, CA, USA

## Abstract

Asgardarchaeota have been proposed as the closest living relatives to eukaryotes, and a total of 72 metagenome-assembled genomes (MAGs) representing six primary lineages in this archaeal phylum have thus far been described. These organisms are predicted to be fermentative organoheterotrophs contributing to carbon cycling in sediment ecosystems. Here, we double the genomic catalogue of Asgardarchaeota by obtaining 71 MAGs from a range of habitats around the globe, including deep subsurface, shallow lake, and geothermal spring sediments. Phylogenomic inferences followed by taxonomic rank normalisation confirmed previously established Asgardarchaeota classes and revealed four novel lineages, two of which were consistently recovered as monophyletic classes. We therefore propose the names *Candidatus* Hodarchaeia class nov. and *Cand.* Jordarchaeia class nov., derived from the gods Hod and Jord in Norse mythology. Metabolic inference suggests that both novel classes represent methylotrophic acetogens, encoding the transfer of methyl groups, such as methylated amines, to coenzyme M with acetate as the end product in remnants of a methanogen-derived core metabolism. This inferred mode of energy conservation is predicted to be enhanced by genetic code expansions, i.e. recoding, allowing the incorporation of the rare 21st and 22nd amino acids selenocysteine (Sec) and pyrrolysine (Pyl). We found Sec recoding in Jordarchaeia and all other Asgardarchaeota classes, which likely benefit from increased catalytic activities of Sec-containing enzymes. Pyl recoding on the other hand is restricted to Hodarchaeia in the Asgardarchaeota, making it the first reported non-methanogenic lineage with an inferred complete Pyl machinery, likely providing this class with an efficient mechanism for methylamine utilisation. Furthermore, we identified enzymes for the biosynthesis of ester-type lipids, characteristic of Bacteria and Eukaryotes, in both novel classes, supporting the hypothesis that mixed ether-ester lipids are a shared feature among Asgardarchaeota.

## Main

The recently described Asgard archaea have been proposed as the closest living prokaryotic relatives to Eukaryotes, supporting a two-domain tree of life [1, 2]. Six Asgard lineages have been described, all of which are named after Norse gods; Lokiarchaeota, Thorarchaeota, Odinarchaeota, Heimdallarchaeota, Helarchaeota, and the recently proposed Gerdarchaeota [1, 3–5]. Asgard archaea were introduced as a superphylum [1], and a subsequent classification, based on taxonomic rank normalisation using relative evolutionary divergence (RED) [6], assigned this lineage to the rank of phylum for which the name Asgardarchaeota with the classes Lokiarchaeia, Thorarchaeia, Odinarchaeia, and Heimdallarchaeia were proposed as Latin placeholder names until nomenclature types are designated [7].

The inferred eukaryotic-like nature of the Asgardarchaeota, in particular the encoded plethora of eukaryotic signature proteins (ESPs), spurred initial speculations about possible eukaryotic contamination of the recovered metagenome-assembled genomes (MAGs)[8]. However, these arguments have since been refuted by analysing additional MAGs [9] and long-read sequencing technologies yielding near-complete MAGs have confirmed that eukaryote-like features are integral to Asgardarchaeota genomes [10]. Furthermore, a recent, decade-long isolation effort resulted in the first Asgardarchaeota isolate, *Candidatus* Prometheoarchaeum syntrophicum strain MK-D1, a Lokiarchaeia representative from deep-sea sediments [11]. The authors obtained a closed genome encoding 80 ESPs and presented evidence for the transcription of these genes, supporting not only that Asgardarchaeota genomes are not chimeric assembly artefacts, but also that ESPs are actively expressed by these archaea.

Insights into the metabolism of Asgardarchaeota based on functions inferred from MAGs, transcriptomics, and experimental data from the Lokiarchaeia culture indicate that members of this phylum are anaerobic fermentative organoheterotrophs [5, 11, 12], although at least some Heimdallarchaeia seem to have acquired oxygen-dependent pathways in their recent evolutionary history [13]. Heimdallarchaeia, Thorarchaeia, and Lokiarchaeia encode the complete archaeal Wood–Ljungdahl pathway [14], which could operate in reverse to oxidize organic substrates and function as an electron sink [12]. It was further hypothesised that cofactors reduced by Asgardarchaeota during organic carbon oxidation may be reoxidized by fermentative hydrogen production to fuel a syntrophic relationship with hydrogen- or formate-consuming organisms [12]. The Lokiarchaeia culture *Ca*. P. syntrophicum MK-D1 confirmed several of these inferred functions. In particular, this archaeon uses small peptides and amino acids while growing syntrophically with a methanogen or a bacterial sulfate reducer through interspecies hydrogen and possibly also formate transfer [11].

Despite this recent focus on Asgardarchaeota, we have likely only explored a small fraction of the diversity encompassed by this phylum. Microbial community profiling based on small subunit (SSU) rRNA gene sequences suggest that many novel Asgardarchaeota lineages are awaiting genomic discovery [5, 14, 15]. Here we describe the recovery of 46 Asgardarchaeota MAGs from coastal, hot spring, and deep-sea sediments complemented by 25 MAGs extracted from public metagenomic datasets. This improved genomic sampling enabled us to resolve phylogenomic relationships, extend the rank normalisation analysis, and to propose two new classes, *Cand*. Hodarchaeia and *Cand*. Jordarchaeia, both named after Norse Gods. Based on metabolic reconstruction we infer both novel lineages to be methylotrophic acetogens, which make use of genetic recodings to enhance their metabolic capabilities.

## Results and discussion

### Sampling sites and community profiling

An *in silico* small subunit (SSU) rRNA gene survey, based on SILVA (r132) [16] revealed 99 sites around the globe, predominantly from anoxic marine and freshwater sediments, as potential Asgardarchaeota habitats for metagenomic recovery (**Fig S1**). Subsequently, we shotgun sequenced and SSU rRNA gene screened local sites in Queensland, Australia, with similar characteristics and discovered Asgardarchaeota in anoxic sediments from two brackish lakes at the Sunshine Coast with relative abundances of up to 2.7% (**Fig. S2a-c**). We extended our search to deep sea sediments and detected Asgardarchaeota in anoxic cores from the Hikurangi Subduction Margin of the Pacific Ocean with relative abundances reaching 11.6% in core segments 1.5 to 634.7 m below the seafloor (mbsf), with the highest abundances reported for depths greater than 100 mbsf (**Fig. S2d,e**). Additionally, we identified two hot spring sediments, from Mammoth Lakes, CA, U.S. and Tengchong, China, as Asgardarchaeota habitats (**Fig. S2f,g)**.

### Genome recovery, phylogenomics, and taxonomic rank normalisation

Metagenomic analysis of the selected lake, deep-sea, and hot spring sediments yielded a set of 46 Asgardarchaeota MAGs, which were supplemented with 25 MAGs recovered from the NCBI Sequence Read Archive (SRA) (**Table S1**). Overall, the 71 MAGs have an average estimated completeness of 78.7±15.3% with an estimated contamination of 3.8±2.3% (**Table S1**). The GC content ranged from 28.8-48.4%, and the average genome size was estimated to be ~4 Mbp (**Table S1; Fig. S3**).

We inferred evolutionary relationships via maximum-likelihood and Bayesian trees (**Table S2**) from trimmed multiple sequence alignments of 122 and 53 archaeal single-copy marker proteins, respectively [17, 18]. Our phylogeny was further evaluated by inferring trees from 1) alignments post removal of compositionally biased sites to increase tree accuracy for distantly related sequences, and 2) alignments of alternative concatenated marker sets including 16 ribosomal proteins (rp1) [19] and 23 ribosomal proteins (rp2) [20]. All phylogenomic inferences of our extended dataset confirmed the monophyly of previously proposed Asgardarchaeota lineages and recovered up to four novel lineages within this phylum (**Fig 1**, **S4-S11**). Next, we applied the taxonomic rank normalisation approach implemented in the Genome Taxonomy Database (GTDB) [6, 7] to assign ranks to Asgardarchaeota lineages. Our results support the rank of class for Thorarchaeia, Odinarchaeia, Heimdallarchaeia, and Lokiarchaeia (**Fig. 1; Table S3**). The previously proposed “Helarchaeota” and “Gerdarchaeota” were robustly placed within these classes and represent the order Helarchaeales in Lokiarchaeia and the order Gerdarchaeales in Heimdallarchaeia, respectively (**Fig. 1; Table S3**). Two of the novel lineages comprised of 4 and 5 MAGs, respectively, were robustly recovered in all phylogenies (**Fig. 1; S4-S11**) and assigned the rank of class based on their RED values and independence from other classes within Asgardarchaeota. A pangenomic analysis based on protein clusters further supported considerable differences between the novel and existing classes (**Fig. S12**). We propose the names *Candidatus* Hodarchaeia class nov. and *Candidatus* Jordarchaeia class nov., derived from the gods Hod and Jord in Norse mythology. We designated type genomes [21] in both lineages (see proposal of type material) and provide a detailed metabolic reconstruction for both classes below. The phylogenetic placement of the remaining two novel lineages, comprising only two MAGs each, from lake and subsurface sediments, respectively (**Table S1; Fig. S2**), was not consistent among trees inferred from different models and marker sets (**Fig. S4-11**), and we therefore assign them the placeholder names, Asgard hot vent group (AHVG) and Asgard Lake Cootharaba group (ALCG). We foresee that the phylogeny of both lineages will be resolved as more Asgardarchaeota genomes become available.

**Figure 1.**
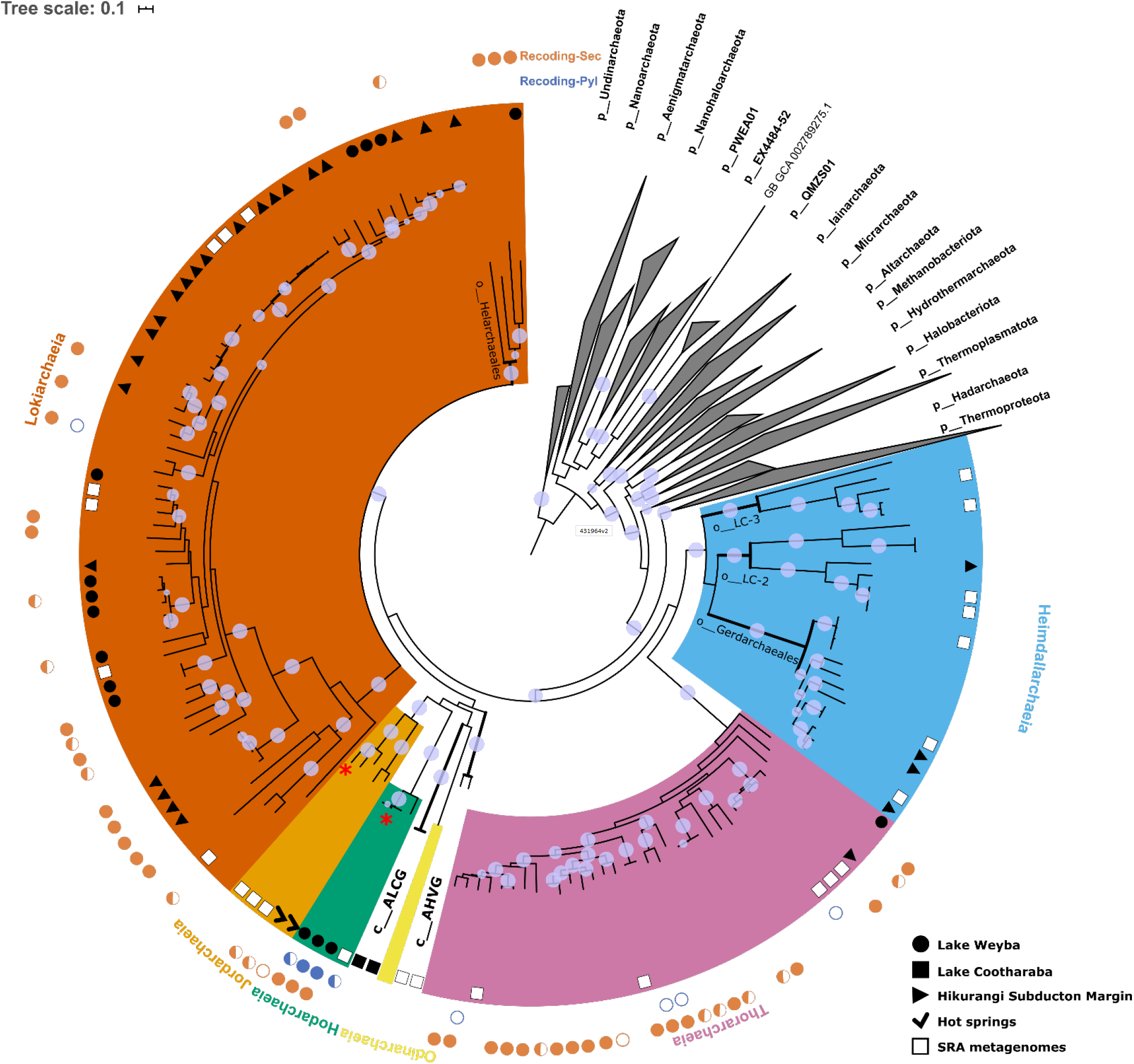
Phylogeny and rank normalised taxonomy of Asgardarchaeota. A maximum-likelihood inference was performed using IQ-TREE under the LG+C10+F+G+PMSF model, based on a multiple-sequence alignment of up to 122 protein markers (subsampled to 42 amino acids per marker) for 143 Asgardarchaeota MAGs, and 1637 archaeal representatives of non-Asgard lineages in GTDB release r95 (1780 taxa, 5124 sites). The tree was rooted on the Undinarchaeota. Branches with bootstrap support >0.9 are depicted with purple circles. Habitat information for Asgardarchaeota MAGs recovered in this study is shown with black and white symbols at the branch tips corresponding to the symbols in the figure legend. The two outer layers indicate the presence of inferred selenocysteine (Sec, orange) and pyrrolysine (Pyl, blue) encoding systems in each MAG: color-filled circles represent a complete Sec/Pyl-encoding system, i.e. genes required for the Sec/Pyl biosynthesis and insertion, and the corresponding tRNA; semicircles represent a partial set of detected genes; empty circles indicate the presence of only a tRNAsec or tRNApyl. Asgardarchaeota classes are highlighted in different colours: Bright cyan - Hodarchaeia; dark yellow - Jordarchaeia; Light pink - Thorarchaeia; orange - Lokiarchaeia; Sky blue - Heimdallarchaeia; Light yellow - Odinarchaeia; Asgard hot vent group (AHVG) and Asgard Lake Cootharaba group (ALCG) - black with bold nodes. Note, that the previously proposed lineages ‘Gerdarchaeota’ and ‘Helarchaeota’ are reassigned to order-level groups and their labels are displayed next to corresponding nodes.

To evaluate the placement of the novel lineages Hodarchaeia and Jordarchaeia in regard to Eukaryotes, we inferred a tree based on 15 markers conserved in the Archaea and Eukaryotes [22]. This inference confirmed previous results by placing Heimdallarchaeia as a sister group to Eukaryotes within Asgardarchaeota, whereas Hodarchaeia and Jordarchaeia clustered with the remaining lineages in this phylum (**Fig. S13**). The detection of numerous eukaryotic signature proteins (ESPs) in Hodarchaeia and Jordarchaeia (**Fig. S14, Table S4**) further supports a close relationship between Asgardarchaeota and eukaryotic organisms. However, the patchy distribution of ESPs in the two novel classes and other Asgardarchaeota lineages (**Fig. S14**), and the observed lack of organelle-like structures in the Lokiarchaeia culture [11], suggests that the ESPs encoded in extant Asgardarchaeota are reminiscent of genes present in the last Asgard archaeal common ancestor (LAsCA) and are likely to perform different functions than their eukaryotic homologs.

### Core metabolism and electron transport

Based on metabolic inference, we propose that Hodarchaeia and Jordarchaeia are methylotrophic acetogens **(Fig. 2, Table S5-S10).** Members of both classes encode enzymes catalysing the transfer of methyl groups, such as methylated amines, to coenzyme M (CoM), similar to a pathway previously reported for methylotrophic methanogens [23]. However, Hodarchaeia and Jordarchaeia are missing genes for methyl-CoM reductase (Mcr), the enzyme catalysing the final step in methane formation. Instead, the encoded tetrahydromethanopterin (H_4_MPT) methyltransferase could facilitate the transfer of methyl groups from methyl-CoM to methyl-H_4_MPT, and subsequently to acetyl-coenzyme A (CoA) to be reduced to acetate for energy conservation (**Fig. 2**). The H_4_MPT methyltransferase might also oxidise the methyl groups via the reverse archaeal Wood-Ljungdahl pathway (WLP; H_4_MPT-dependent). Alternatively the WLP could function in the opposite direction to autotrophically fix carbon dioxide using hydrogen as an electron donor, however we did not detect genes of uptake hydrogenases, i.e. Hodarchaeia and Jordarchaeia lack genes for group 1 NiFe-hydrogenases (**Table S10**).

**Figure 2.**
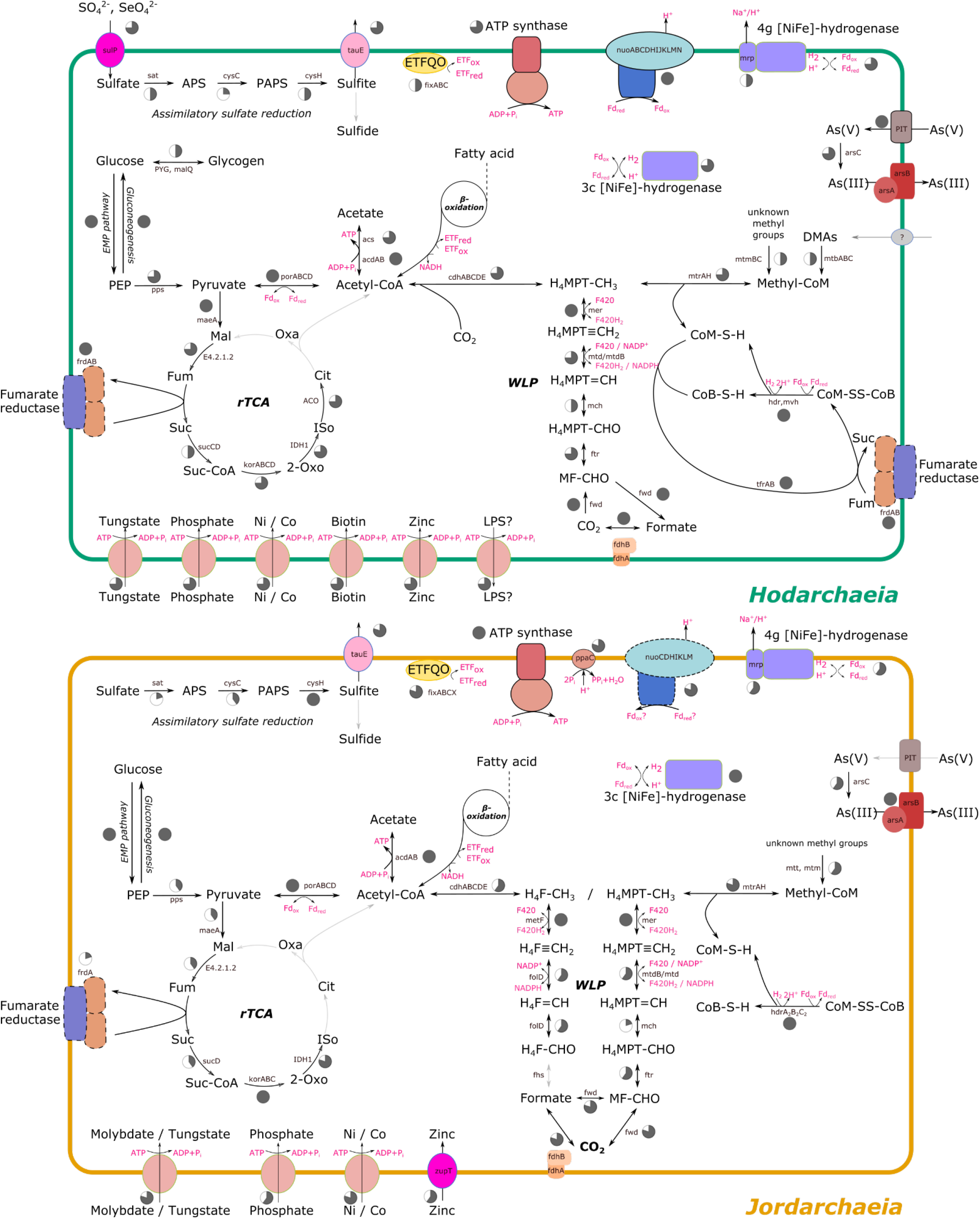
Inferred metabolism of Hodarchaeia and Jordarchaeia. Each of the arrows represents functions assigned to predicted proteins encoded in the respective genomes (Hodarchaeia, 4 MAGs, max. completeness 94.1%); Jordarchaeia, 5 MAGs, max. completeness 95.0%). Pie charts next to the arrows/enzymes indicate the proportion of MAGs that encode a certain enzyme (See Table S4-S10 for more information). A black arrow indicates that this enzymatic step is encoded by at least one MAG of a given class (Hodarchaeia or Jordarchaeia), and a solid grey arrow indicates an enzymatic step that is missing from a given class. The presence or function of enzyme complexes surrounded by the black dashed lines is tentative. Note that Hodarchaeia and Jordarchaeia genomes encode an acetyl-CoA synthetase (acs) which preferentially acts in the direction of acetate and ATP production. ***Abbreviations***: APS, adenylyl sulfate; PAPS, 3’-Phosphoadenylyl sulfate; PEP, phosphoenolpyruvate; H4F, tetrahydrofolate; H_4_MPT, tetrahydromethanopterin; DMA, dimethylamine; MMA, trimethylamine; LPS, lipopolysaccharide; CoM, coenzyme M; Oxa, oxaloacetate; Cit, citrate; Iso, isocitrate; 2-Oxo, 2-oxo-glutarate; Suc-CoA, succinyl-coenzyme A; Suc, succinate; Fum, fumarate; Mal, malate; ETF, electron transfer flavoprotein; ETFQO, ETF-ubiquinone oxidoreductase; Fd, ferredoxin; Pi, inorganic phosphate; PPi, inorganic pyrophosphate; red, redox; rTCA, reverse tricarboxylic acid cycle; WLP, Wood-Ljungdahl pathway.

The genomes of both novel classes encode the ability to break down complex carbohydrates via glycoside hydrolases including β-galactosidase and α-amylase, and carbohydrate esterases (**Table S8**). The resulting glucose could be utilised via the Embden–Meyerhof–Parnas (EMP) pathway to generate pyruvate, for the subsequent oxidation to acetate, or to be metabolised by the encoded partial reverse TCA cycle (**Fig. 2**). Electrons derived from oxidizing these organic substrates and various methyl groups could establish a membrane potential since Hodarchaeia and Jordarchaeia encode genes for complex I (dehydrogenase) and complex V (ATP synthase) of the electron transport chain (**Fig. 2).** Notably, complex I lacks the reduced cofactor oxidizing subunits NuoEFG, which form the NADH dehydrogenase module, therefore we hypothesise that energy conservation in Hodarchaeia and Jordarchaeia depends on electron transfer by reduced ferredoxin (**Fig. S15**), similar to the membrane-bound fpo-like complex of the acetoclastic methanogens [24]. Oxidised ferredoxin could be reduced by the encoded cytoplasmic heterodisulfide reductase [NiFe]-hydrogenase complex while oxidising CoM-CoB heterodisulfide to coenzyme B and coenzyme M (**Fig. 2, S15).** In Hodarchaeia, both coenzymes might be re-oxidised by the encoded thiol:fumarate reductase which catalyses the reduction of fumarate, with CoB and CoM as electron donors, to succinate and heterodisulfide CoM [25]. Alternatively electrons could be shuttled to a syntrophic partner via extracellular electron transfer mediated by hydrogen, formate, or acetate [26]. A symbiotic relationship would also help to complement the amino acid needs of Hodarchaeia and Jordarchaeia, which lack genes encoding the biosynthesis of the amino acids proline, tyrosine, and phenylalanine, and additionally alanine biosynthesis genes are missing in Jordarchaeia (**Table S9**).

### Mixed membrane lipids and the great lipid divide

Both Hodarchaeia and Jordarchaeia encode all genes for the synthesis of archaeal ether-type lipids, but in addition, Hodarchaeia also encode enzymes for the biosynthesis of ester-type lipids, characteristic of Bacteria and Eukaryotes (**Fig. S16**). This finding aligns with previous reports of ester lipid biosynthetic pathways in Asgard lineages [27], supporting the hypothesis that mixed ether-ester lipids are a shared feature among Asgardarchaeota. Subsequently, this trait could have been lost in some subordinate lineages, including Jordarchaeia (**Fig. S16**). Phylogenetic inference of a key ester-type lipid gene supports the finding that archaeal homologs are distinct from their bacterial counterparts [28] and showed some Lokiarchaeia genes clustering with eukaryotic homologs, albeit with low support values (**Fig. S17**). The great lipid divide between bacteria and archaea has been further eroded by the discovery of ester-type lipid genes in members of the Poseidoniales (Marine Group II archaea) [29], and functional validation of ether-type lipid genes in the Fibrobacteres–Chlorobi–Bacteroidetes (FCB) superphylum [30]. This suggests, together with the reported extensive interdomain horizontal gene transfer of several membrane lipid biosynthesis genes [28], that the lipid divide thought to distinguish the domains of life might be more permeable than previously thought.

### Transporters

Both Hodarchaeia and Jordarchaeia encode several ABC transporters for the uptake of essential trace compounds, including tungstate [31], which has been shown to enhance the growth of methanogens [32] and could provide a similar benefit to both classes (**Fig. 2, S18; Table S9)**. In addition, Hodarchaeia possess a low-affinity inorganic phosphate transporter that also functions as a major uptake system for arsenate [33]. To mitigate the toxicity of arsenate, both classes may be able to actively expel arsenate from their cells by reducing arsenate to the less toxic arsenite [34], which can then be pumped out of the cell by the ATP-consuming arsenite exporter (**Fig. 2)**.

### Expanding metabolic capabilities by recoding the genetic code

Based on inferred proteins and codon usage we predict that Hod and Jordarchaeia increase their amino acid repertoire and consequently their metabolic potential through localised recoding strategies. These include the recoding of the STOP codons opal (UGA) and amber (UAG) to incorporate the rare 21st and 22nd amino acid selenocysteine (Sec) and pyrrolysine (Pyl), through distinct recoding processes. Both novel classes encode the archaeal/eukaryotic-type Sec biosynthesis machinery. In particular, we detected a single selenocysteine t-RNA (tRNAsec) in Hodarchaeia and Jordarchaeia MAGs (**Fig. 3 c,d**) and confirmed previous reports of this tRNA in Lokiarchaeia and Thorarchaeia [35, 36], but did not identify a tRNAsec in Heimdallarchaeia or Odinarchaeia (**Table S11**). Remarkably, the tRNAsec in all Hodarchaeia and some Lokiarchaeia had unusual insertions and deletions, negating previously proposed domain-specific characteristics. For example, the Hodarchaeia tRNAsec has a short 6 bp D-stem **(Fig. 3, S19**), a feature that has been attributed to eukaryotes and bacteria, whereas archaeal tRNAsec were thought to generally possess a 7 bp D-stem [37]. Our tRNAsec phylogeny recovered most recoded Asgardarchaeota lineages as monophyletic groups clustering with methanogens [37] and Eukaryotes, albeit with low bootstrap support values likely due to the short alignment length (**Fig. S20**). The recovery of monophyletic tRNAsec groups that match the species tree suggest that horizontal gene transfers (HGTs) may not be common in the evolutionary history of tRNAsec, despite the reported frequent and extensive gene duplication of tRNAs in general [38]. Furthermore, Hodarchaeia and Jordarcheia, as well as Lokiarchaeia and Thorarchaeia, encode enzymes to correctly charge this tRNA in order to synthesise a functional selenocysteine tRNAsec (Sec-tRNAsec) using the archaeal/eukaryotic-type Sec biosynthesis pathway. This process involves an initial mischarging of tRNAsec with serine by seryl-tRNA synthetase, then phosphorylation by phosphoseryl-tRNA kinase and conversion into a functional tRNAsec by Sec synthase using selenophosphate formed by selenophosphate synthetase (SPS) from selenium **(Fig. 3a**) [39]. We found no evidence for the presence of a bacterial-type Sec biosynthesis pathway in Asgardarchaeota, despite previous reports of a bacterial Sec synthase (SelA) (**Fig. 3a**) in Thorarchaeia MAG SMTZ1-83 [36]. Instead, we suggest that the contig harbouring SelA in this MAG is likely bacterial contamination (**Table S12**), leading us to posit that Asgardarchaeota rely solely on selenophosphate-dependent synthesis of Sec-tRNASec (**Fig. 3a**).

**Figure 3.**
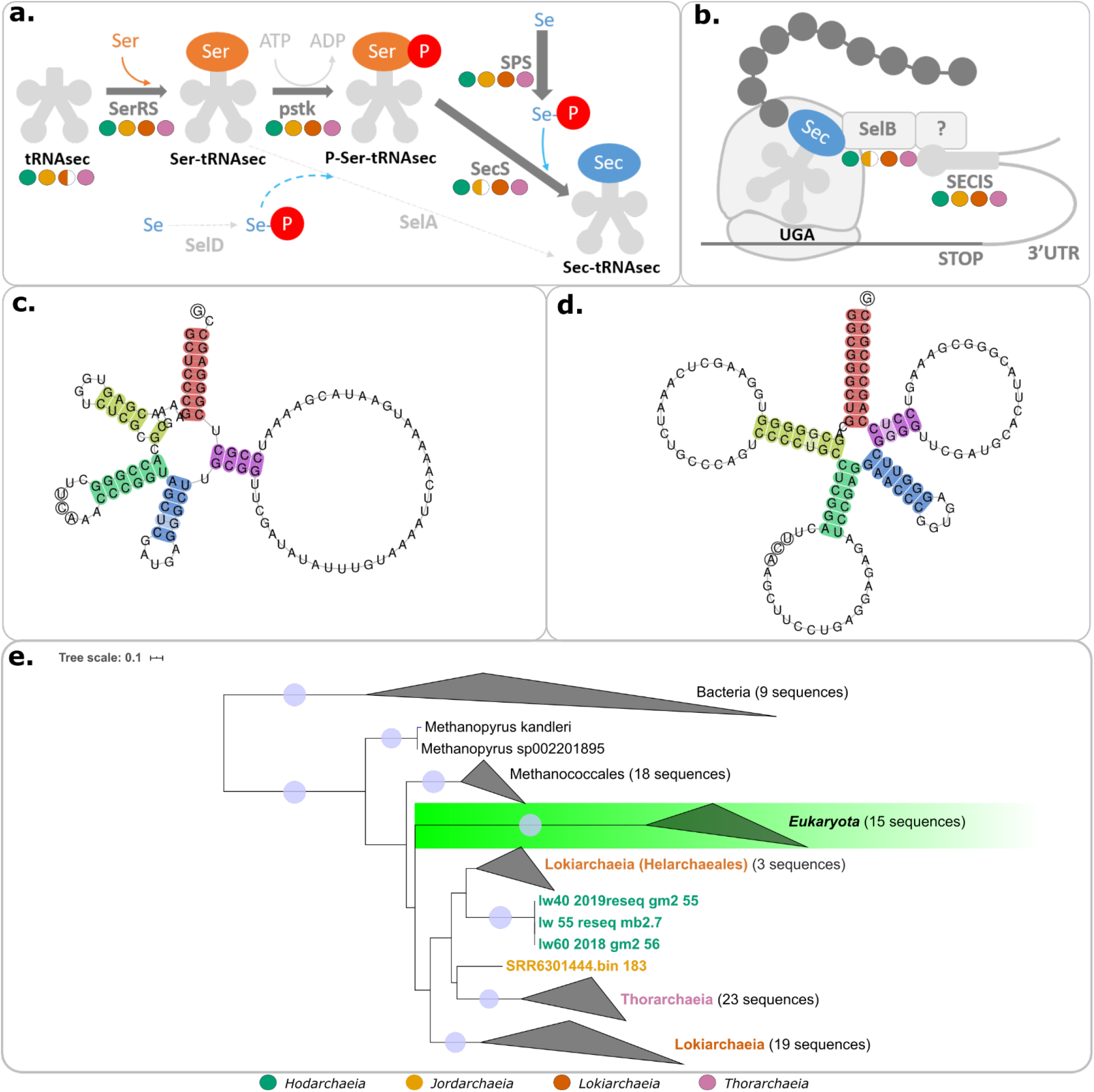
Selenocysteine recoding in Asgardarchaeota. **a.** Proposed mechanism of selenocysteine biosynthesis in Asgardarchaeota. **b.** Proposed selenocysteine insertion in Asgardarchaeota. Presence of genes in each class are indicated with coloured circles: filled circle - half or more of the MAGs encode the given gene; half-filled circle - less than half of the MAGs encode the given gene. Note that the eukaryotic SECIS-binding protein 2 (SBP2) is missing from all Archaea (indicated by a question mark). **c.** tRNAsec in Hodarchaeia MAG “lw60_2018_gm2_56” and **d.** tRNAsec in Jordarchaeia MAG “1308”. Highlighted are the acceptor arm (red), D arm (yellow), anticodon arm (green), variable arm (blue), and the T arm (purple). **e.** Phylogenetic tree of *selB*, inferred with IQ-TREE (PMSF C10 model) from a TrimAl-trimmed alignment of SelB genes from archaeal, bacterial, and eukaryotic genomes. Tree was rooted between the bacterial and archaeal-eukaryotic clade. Asgardarchaeota sequences are highlighted with different colour labels: Bright cyan - Hodarchaeia; dark yellow - Jordarchaeia; Light pink - Thorarchaeia; orange - Lokiarchaeia.

Sec insertion in Hodarchaeia and Jordarcheia and all other Sec recoded Asgardarchaeota lineages could be mediated by the Sec-specific elongation factor (SelB), which connects the selenocysteine insertion sequences (SECIS), an RNA element that forms a stem-loop structure during Sec insertion (**Fig. 3b**), to the ribosome with the help of the SECIS-binding protein 2 (SBP2). Phylogenetic analysis of SelB and SPS supports a predominantly vertical inheritance of both genes and a separation of bacterial and archaeal/eukaryal orthologs (**Fig. 3e**, **S21-22**). Within the archaeal/eukaryal branch, the genus *Methanopyrus* was identified as the deepest branching lineage in both trees, and Asgardarchaeota formed a monophyletic sister group to Eukaryotes, although with low bootstrap support. We did not detect SBP2 homologs in Asgardarchaeota, consistent with previous reports that Archaea do not encode this elongation factor, and implying that this key enzyme evolved after eukaryogenesis [35, 39]. We found 12 to 25 and 19 to25 predicted SECIS elements, the site where Sec insertion occurs, in Hodarchaeia and Jordarchaeia MAGs, and three proteins that include a selenocysteine, i.e. selenoproteins (**Table S13**). The detected selenoproteins were located 30 to 500 bases upstream of the corresponding SECIS element, a distance range previously observed in Archaea and Eukaryotes [35, 40]. All three selenoproteins detected in Hodarchaeia and Jordarchaeia, including the F_420_-non-reducing hydrogenase (Vhu) subunit D and the heterodisulfide reductase (Hdr) subunit A, are also present in Lokiarchaeia [35]. This suggests that selenoproteins are common across all Asgardarchaeota, which likely depend on the increased catalytic activity of Sec-containing proteins, such as HdrA and Vhu, as part of their energy conservation strategies (**Fig. S15**). Indeed, it has been experimentally verified that selenoproteins can provide up to a hundred times increased catalytic activity over cysteine, its sulphur-containing analogue [41]. Further support for a Sec-enhanced metabolism among Asgardarcheota are sulfate permeases (sulP), which are encoded in three Hodarchaeia and several Lokiarchaeia genomes, and could import sulfate and related oxyanions such as selenate, the oxidised form of selenium [42–44].

### Pyl recoding

We detected a second recoding solely present in Hodarchaeia which affects the amber (UAG) stop codon and could allow this class to use the rare 22nd amino acid pyrrolysine (Pyl). The presence of a Pyl tRNA, all required Pyl biosynthesis genes, and specific Pyl-encoded proteins suggests that this recoding provides Hodarchaeia with an efficient mechanism for methylamine utilisation, despite an unusually high UAG stop codon usage.

Hodarchaeia encode a complete Pyl encoding system including all three Pyl biosynthesis proteins (PylB, PylC, PylD) and a pyrrolysyl-tRNA synthetase (PylS) to charge the pyrrolysine tRNA (tRNApyl, pylT) (**Fig. 4a**) [45]. Unlike selenocysteine (**Fig. 3b**), no specific proteins or insertion sequences are required for the tRNApyl insertion, which has been proposed to directly compete with the translation termination release factor for UAG codons (**Fig. 4b**) [46]. While Pyl genes in Archaea usually form an uninterrupted pylTSBCD cluster, Hodarchaeia show a pattern similar to *Methanohalobium evestigatum* [47], in which the pylS gene is ~6 Kb distant from pylBCD, separated by a NAD kinase and several hypothetical proteins (**Fig. 4c**). The tRNApyl of Hodarchaeia, encoded by pylT, is located upstream of pylS and displays a classic cloverleaf secondary structure with an unusual acceptor stem tail that discriminates the Hodarchaeia tRNApyl from the CCA tails of previously reported archaeal and bacterial homologs (**Fig. 4d**) [48]. Remarkably, the reported low usage (<6%) of the UAG STOP codon in Pyl-containing Archaea [46, 49] does not apply to Hodarchaeia, with 27% of their CDSs terminating with UGA (**Fig. 4e**; **Table S14**), a percentage corresponding to UGA frequencies of Pyl-encoding bacteria [50]. How a mis-specification of Pyl-tRNApyl to the frequent UAG stop sense codons is avoided remains unknown, although possible mechanisms exist (see below).

**Figure 4.**
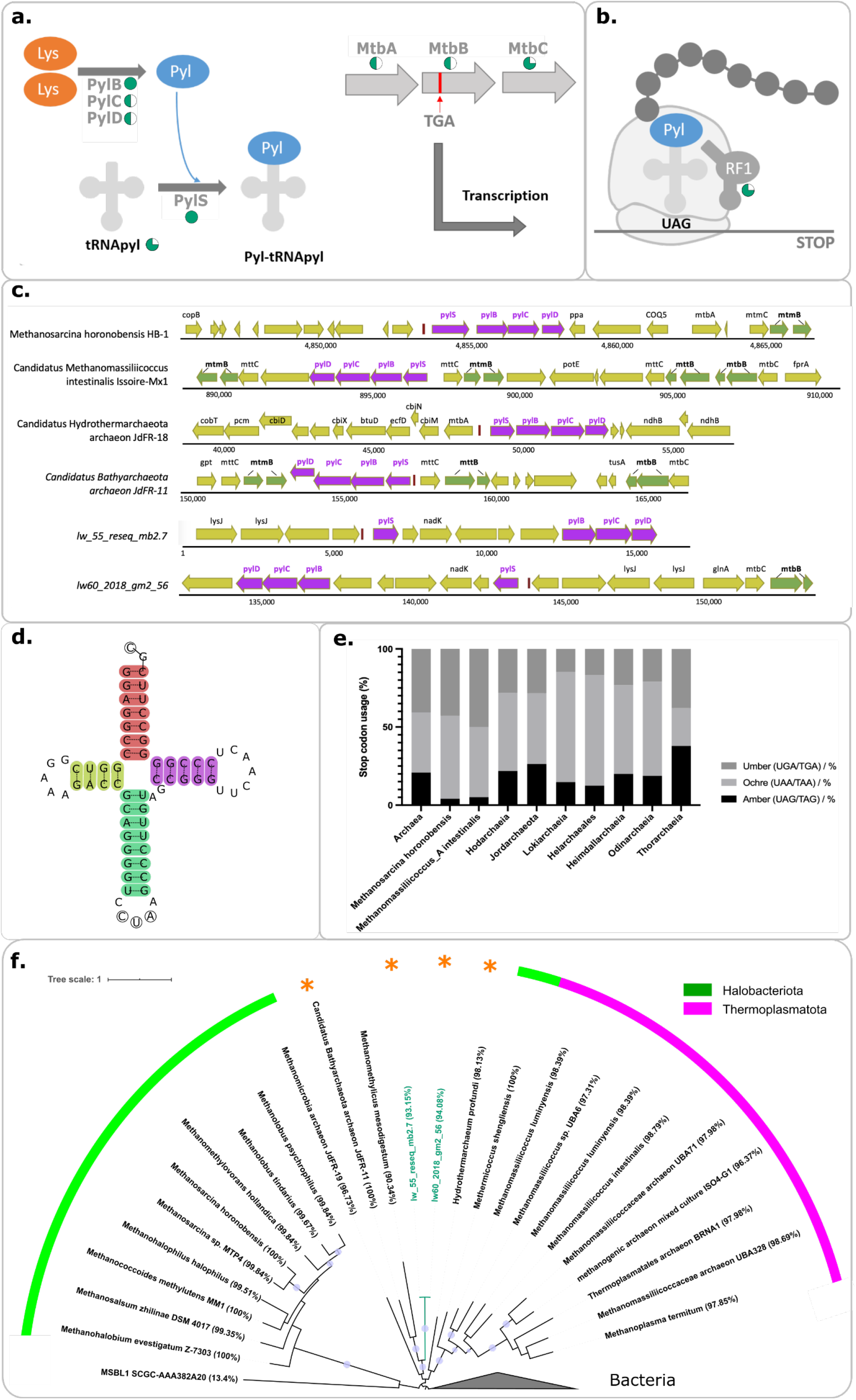
Pyrrolysine recoding machinery and stop codon usage. **a.** Proposed pyrrolysine (Pyl) biosynthesis in Hodarchaeia. **b.** Proposed Pyl insertion in Hodarchaeia. The proportion of Hodarchaeia MAGs bearing Pyl-recoding machinery genes is indicated with bright cyan pie charts. **c.** Gene neighbourhood of the Pyl cluster. Gene names are labelled above the corresponding CDS. Pyl cluster genes (pylSBCD) are highlighted in dark pink, and pyrrolysine-containing genes are highlighted in green. **d.** tRNApyl in Hodarchaeia. The highlighted regions are the acceptor arm (red), the D arm (yellow), the CUA anticodon arm (green), and the T arm (purple). The acceptor stem in Hodarchaeia displayed a GC tail, which is distinct from previously reported archaeal and bacterial tRNApyl which have a CCA tail. **e.** Stop codon usage in Asgardarchaeota, and two recoded lineages of methanogens. **f.** Maximum-likelihood tree (IQtree with 100 bootstraps replicates) based on a concatenated alignment of pylSBCD genes. Purple circles represent branches with bootstrap support over 0.9. The two Hodarchaeia sequences are highlighted with cyan branches and labels. Genome completeness values calculated by CheckM are provided in brackets after each organism name. All taxa for which we report a Pyl cluster for the first time, i.e. *Candidatus* Bathyarchaeota archaeon JdFR-11, *Candidatus* Hydrothermarchaeum profundi, and Hodarchaeia are indicated with an orange asterisk. See **Fig. S24** for a rooted PylSBCD tree and **Fig. S25-28** for individual gene trees.

Pyl recoding has only been reported previously in archaeal methanogens belonging to the archaeal phyla Thermoplasmatota and Halobacteriota, the class Methanomethylicia (Verstraetearchaeota, *sensu* NCBI taxonomy), and from the candidate lineage *Persephonarchaea MSBL1* [49, 51, 52]. Several bacterial phyla, including Firmicutes and Desulfobacterota, also possess Pyl recoding thought to be acquired from Archaea via multiple HGTs [53], but this recoding is absent in eukaryotes [48]. In addition to Hodarchaeia, we identified pylSBCD genes for the first time in Hydrothermarchaeota and Bathyarchaeia representatives by mining GTDB genomes [6], and most gene phylogenies support a novel cluster containing both representatives, together with Methanomethylicia, and Hodarchaeia (**Fig. 3f**, **S24-28**).

The major role of Pyl recoding in Archaea, methanogenic and non-methanogenic alike, is methylamine utilisation, since Pyl is foremost incorporated in the active sites of methyltransferases [49, 54]. Indeed, Hodarchaeia encode several methyltransferases, including monomethylamine methyltransferase (MtmB) and dimethylamine methyltransferase (MtbB), with the latter possessing a Pyl recoding, making it the only in-frame UAG stop codon in Hodarchaeia (**Table S15; Fig. 4c**). Thereby, MtbB, together with Methylcobamide:CoM methyltransferase (MtbA), could methylate the cognate corrinoid protein (MtbC), which in turn methylates coenzyme M (CoM) [47]. This cascade of encoded methyl transfers should allow Hodarchaeia to convert methyl groups directly to acetate for energy conservation (see above). Hence, maintaining the Pyl-recoding seems essential, since MtbB requires Pyl, which was hypothesized to activate and orient methylamines as substrates for the corrinoid protein MtbC [47]. How Hodarchaeia, with their high percentage of UAG stop codons, control the specificity of Pyl insertions versus protein termination remains to be determined, however, it has been suggested that environmental conditions such as the presence of methylamines could selectively activate Pyl biosynthesis [46]. Indeed, the Firmicute *Acetohalobium arabaticuma* was recently found to expand its genetic code to include Pyl only in the presence of trimethylamines (TMA), but to down-regulate the transcription of the entire Pyl operon when TMA was absent[50].

### Recoding evolutionary history in Asgardarchaeota

While the evolutionary history of Pyl encoding is still debated, a structure-based phylogeny suggested that PylS was present in the last universal common ancestor (LUCA) [53, 55]. Similarly, it has been argued that Sec recoding is an ancient trait considering the highly conserved nature of the Sec incorporation machinery [35], and the fact that the genes involved are not always physically linked in an operon, which impedes its propagation between lineages via horizontal transfer [56]. Our Pyl and Sec trees indicate primarily vertical evolution of these genes (**Fig. S20-28**), suggesting that HGT is an infrequent event in the evolution of both traits in archaea. Therefore, we suggest that the last Asgard archaeal common ancestor (LAsCA) possessed both the Pyl and Sec recoding. Subsequently, Pyl was lost in the branch leading to Heimdallarchaeia and Eukaryotes, and also in Jordarchaeia and Odinarchaeia (**Fig. 5**). Lokiarchaeia and Thorarchaeia also lack the Pyl gene cluster (**Fig. 5; Fig. 1; Table S15**), but we detected tRNApyl sequences in genomes from both lineages which could be remnants of an ancient Pyl trait that has since been lost. The roles of these tRNAs are unknown, however, they could function as sources of various small noncoding RNA species [57]. Sec recoding, on the other hand, remained present in most Asgardarchaeota lineages and was only lost in Heimdallarchaeia and Odinarchaeia (**Fig. 5).** Maintaining these presumably ancient recodings could be driven by selective metabolic advantages, i.e. the catalytic advantages of Sec containing enzymes and the importance of Pyl for active sites of methyltransferases (see above).

**Figure 5.**
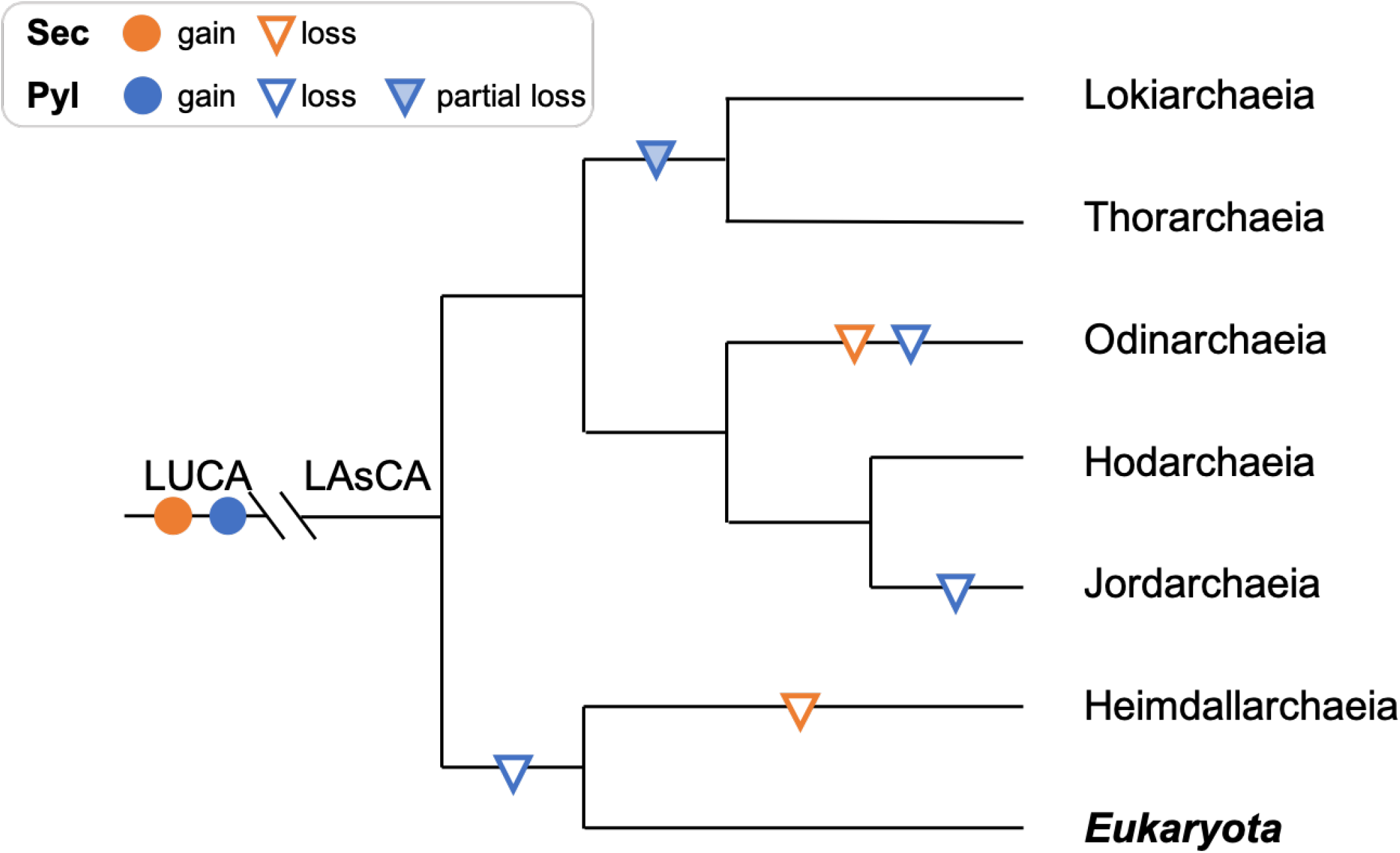
Proposed evolutionary history of pyrrolysine and selenocysteine recoding in Asgardarchaeota. Cladogram, based on the maximum-likelihood tree of 15 ribosomal protein markers (**Fig. S13**), showing gain and loss of selenocysteine (Sec) and pyrrolysine (Pyl) recoding form to the last Asgard archaeal common ancestor (LAsCA) to extend taxa in the Asgardarchaeota including eukaryotes. Partial loss is defined as the loss of Pyl biosynthesis and insertion genes while retaining the Pyl-tRNA. Acronyms: last universal common ancestor (LUCA).

### Conclusion

In the present study, we apply taxonomic rank normalisation to genome phylogenies including 71 novel Asgardarchaeota genomes and propose two novel classes, Hodarchaeia and Jordarchaeia, which have the potential to convert C1 compounds into organic products as methylotrophic acetogens. Thereby, both classes utilise a methanogen-like pathway but do not encode homologs of the key enzyme methyl-CoM reductase (Mcr). This absence, together with the inferred Mcr-like enzymes in Helarchaeales [4], and our detection of an mcrA-like gene in Lokiarchaeia outside the order Helarchaeales (**Fig. S29**), suggests that pathways for the utilisation of methane and other hydrocarbon gases, or remnants thereof, played an important role in the evolution of Asgardarchaeota. We further reveal recoding as an ancient trait in this phylum, which allows the incorporation of the rare amino acids selenocysteine (Sec) and pyrrolysine (Pyl) into selected proteins, possibly yielding benefits from enhanced catalytic properties of Sec and Pyl containing enzymes. Thereby, Pyl, which is restricted to Hodarchaeia, with remnant tRNAs in Thor and Lokiarchaeia, likely supports efficient methylamine utilisation, and possibly represents another relic from a methylotrophic methanogenic or methanotrophic ancestor. Our results support previous reports of a close relationship between Asgardarchaeota and Eukaryotes, based on phylogenetic inferences, the detection of various encoded eukaryotic signature proteins and of enzymes for the biosynthesis of bacteria/eukaryotic-type ester lipids in Hodarchaeia and Jordarchaeia and other lineages in this phylum. We envision that future recoveries of additional Asgardarchaeota MAGs, in concert with culture-based approaches, will further fuel phylogenomic and metabolic reconstructions and lead to the experimental verification of encoded functions, thereby ultimately shedding more light on the origin of Eukaryotes.

### Proposal of type material

#### *Candidatus* Hodarchaeum gen. nov

*Candidatus* Hodarchaeum (Hod.ar.chae’um. N.L. neut. n. *archaeum* archaeon; N.L. neut. n. *Hodarchaeum* an archaeon named after Hod, a blind warrior god in North mythology). Inferred to be a methylotrophic heterotroph with a genetic code expansions, i.e. recoding, allowing the incorporation of the rare 21st and 22nd amino acids selenocysteine and pyrrolysine. Type species: *Candidatus* Hodarchaeum weybense.

#### *Candidatus* Hodarchaeum weybense

*Candidatus* Hodarchaeum weybense (wey.ben’se. N.L. neut. adj. *weybense* of or pertaining to Lake Weyba, a saltwater lake in Queensland, Australia). This uncultured lineage is represented by the genome “lw60_2018_gm2_56”, NCBI WGS **XXXXXX**, recovered from Lake Weyba sediments, and defined as high-quality draft MAG[58] with an estimated completeness of 94.08% and 3.74% contamination, the presence of a 23S, 16S, and 5S rRNA gene and at 16 tRNAs.

#### *Candidatus* Jordarchaeum gen. nov

*Candidatus* Jordarchaeum (Jord.ar.chae’um. N.L. neut. n. *archaeum* archaeon; N.L. neut. n. *Jordarchaeum* an archaeon named after Jord, the goddess of the earth in North mytholody). Inferred to be a methylotrophic heterotroph with a genetic code expansions, i.e. recoding, allowing the incorporation of the rare 21st amino acid selenocysteine. Type species: *Candidatus* Jordarchaeum madagascariense.

#### *Candidatus* Jordarchaeum madagascariense

*Candidatus* Jordarchaeum madagascariense (ma.da.ga.scar.i.en’se. N.L. neut. adj. *madagascariense* of or pertaining to Madagascar, an island country in the Indian Ocean). This uncultured lineage is represented by the genome “A_PRJDB_7”, NCBI WGS **XXXXXX**, recovered from elephant bird fossils in Madagascar, with an estimated completeness of 95.02% and a contamination of 2.41%, the presence of a 23S, 16S, and 5S rRNA gene and at 6 tRNAs.

### Descriptions of higher taxonomic ranks

#### *Candidatus* Hodarchaeaceae fam. nov

Description of Ca. Hodarchaeaceae fam. nov. (Hod.ar.chae. ace’ae. N.L. neut. n. *Hodarchaeum*, Candidatus generic name; -*aceae*, ending to designate a family; N.L. fem. pl. n. *Hodarchaeaceae*, the *Hodarchaeum* family). The family is circumscribed based on concatenated protein phylogeny and rank normalisation approach as per Parks et al., (2018). Type genus is *Candidatus* Hodarchaeum. The description is the same as for *Candidatus* Hodarchaeum gen. nov.

#### *Candidatus* Jordarchaeaceae fam .nov

Description of Ca. Jordarchaeaceae fam .nov. (Jord.ar.chae. ace’ae. N.L. neut. n. *Jordarchaeum*, Candidatus generic name; -*aceae*, ending to designate a family; N.L. fem. pl. n. *Jordarchaeaceae*, the *Jordarchaeum* family. Type genus is *Candidatus* Jordarchaeum. The description is the same as for *Candidatus* Jordarchaeum gen. nov

#### *Candidatus* Hodarchaeales ord. nov

Description of Ca. Hodarchaeales ord. nov. (Hod.ar.chae. a’les. N.L. neut. n. *Hodarchaeum*, Candidatus generic name; -*ales*, ending to designate an order; N.L. fem. pl. n. *Hodarchaeales*, the *Hodarchaeum* order). Type genus is *Candidatus* Hodarchaeum. The description is the same as for *Candidatus* Hodarchaeum gen. nov.

#### *Candidatus* Jordarchaeales ord .nov

Description of Ca. Jordarchaeales ord. nov. (Jord.ar.chae. a’les. N.L. neut. n. *Jordarchaeum*, Candidatus generic name; -*ales*, ending to designate an order; N.L. fem. pl. n. *Jordarchaeaceae*, the *Jordarchaeum* order. The order is circumscribed based on concatenated protein phylogeny and rank normalisation approach as per Parks et al., (2018). Type genus is *Candidatus* Jordarchaeum. The description is the same as for *Candidatus* Jordarchaeum gen. nov

#### *Candidatus* Hodarchaeia class. nov

Description of Ca. **Hodarchaeia class. nov.** (Hod.ar.chae. ia. N.L. neut. n. *Hodarchaeum*, Candidatus generic name; -*ia*, ending to designate a class; N.L. neut. pl. n. *Hodarchaeia*, the *Hodarchaeum* class). Type order is *Candidatus* Hodarchaeales. The description is the same as for *Candidatus* Hodarchaeum gen. nov.

#### *Candidatus* Jordarchaeia class. nov

Description of Ca. **Jordarchaeia class. nov.** (Jord.ar.chae. ia. N.L. neut. n. *Jordarchaeum*, Candidatus generic name; -*ia*, ending to designate a class; N.L. neut. pl. n. *Jordarchaia*, the *Jordarchaeum* class. Type order is *Candidatus* Jordarchaeales. The description is the same as for *Candidatus* Jordarchaeum gen. nov

## Methods

### Small subunit (SSU) rRNA gene *in silico* survey

The small subunit (SSU) rRNA gene survey was based on the SILVA SSU database (release 132, Ref NR 99) [16] (https://www.arb-silva.de/). We extracted the habitat information (field ‘habitat_slv’, ‘isolation_source’ and ‘lat_lon’ in SILVA ARB database) and manually removed habitat entries whose details are duplicated or ambiguous. The remainder of the habitat entries were grouped into seven categories: ‘sediment marine’, ‘sediment freshwater’, ‘sediment other’, ‘microbial mats/biofilms’, ‘soil/permafrost’, and ‘other’’.

### Sample collection and DNA extraction

#### Sunshine Coast Lakes sediment

Lake sediment samples from Lake Cootharaba (LC) (−26.28°, 152.99°) and Lake Weyba (LW) (−26.44°, 153.06°) were sampled using sterilised one-meter PVC pipes. LC sediments at depths from 5 cm to 25 cm and LW sediments at depths from 5 cm to 60 cm were sampled in 5 cm intervals in December 2018 and November 2019. Salinity of lake water was recorded using a Seawater Digital Refractometer (Milwaukee, US). Collected sediments were flash frozen in alcohol and dry ice, and delivered to ALS Environmental testing, Brisbane, Australia for chemical analysis. DNA was extracted within four hours of sampling using the PowerSoil DNA Isolation kit (MoBio, USA) following the manufacturer’s protocol.

#### Hikurangi Subduction Margin sediment

Deep-sea sediment samples of Hikurangi Subduction Margin were sampled by the International Ocean Discovery Program (IODP) Expedition 375 scientists onboard the [59]. Sampling holes were drilled at four sites: U1518 (an active fault near the deformation front; sampling depths range from 0 mbsf to 494.90 mbsf), U1519 (the upper plate above the high-slip slow slip event source region; sampling depths range from 0 mbsf to 640.00 mbsf), U1520 (the incoming sedimentary succession in the Hikurangi Trough; sampling depths range from 0 mbsf to 1045.75 mbsf), and U1526 (atop the Tūranganui Knoll Seamount; sampling depths range from 0m to 83.60 mbsf) [60]). Sediment cores were sub-sampled shipboard using 5ml syringes, which were stored and shipped on dry ice until they reached the laboratory and were then stored at −80°C until DNA extraction. To minimise possible contamination, we trimmed off the outer centimetre of each sample and used the inner sediment core for DNA extraction. To optimise DNA extraction for these low biomass samples, 300 mg sediments were first mixed with G2 DNA/RNA Enhancer beads (Ampliqon, Denmark). The subsequent DNA extraction steps were conducted using the PowerSoil DNA Isolation kit (MoBio, USA) following the manufacturer’s protocol.

#### Geothermal spring sediments

Geothermal spring sediments (top 1 cm) were collected from Little Hot Creek, near Mammoth Lakes, CA, USA, from LHC4 (N37°41.436’, W118°50.653’; 81.1°C; pH=6.83) and Jinze Pool located in Dientan, Tengchong County, China (N23.44138°, E98.46004°; 78.2°C; pH=6.65). Subsamples were stored and shipped on dry ice until they reached the laboratory and were then stored at −80°C till DNA extraction. DNA was then extracted from freshly thawed sediment samples using the FastDNA™ SPIN Kit for Soil (MP Biomedicals, Santa Ana, CA) following the manufacturer’s protocol. The physicochemical conditions in Little Hot Creek (LHC4) and Jinze Pool are described in detail elsewhere [61, 62].

### Shotgun sequencing

For the Hikurangi Subduction Margin and Sunshine Coast lake samples Illumina Nextera XT libraries were constructed and shotgun sequenced using NextSeq 500/550 High Output v2 2 x 150bp paired end chemistry. For the geothermal spring sediments Truseq short-insert paired-end libraries were constructed with an average insert size of 270bp and sequenced on the Illumina HiSeq 2000/2500 1T platform.

### Public data acquisition

Potential Asgardarchaeota containing metagenomes were identified in the NCBI Sequence Read Archive (SRA) using SingleM (https://github.com/wwood/singlem). This software uses single copy marker genes to search for public metagenomes containing reads that match a bacterial or archaeal lineage of interest. The search for Asgardarcheota reads yielded matches for seven corresponding study IDs (SRP029382, SRP061771, ERP013176, SRP077065, SRP049601, DRP003377, and SRP098167) in the SRA database (**Table S1**). Information from all NCBI sequencing runs from each study was collected, but only shotgun metagenomic sequence runs were downloaded for our analysis.

### Small subunit (SSU) rRNA gene community profiles

To obtain microbial community profiles, we aligned the reads of all shotgun sequenced samples to the SILVA 132_99 database [16] and classified the reads into operational taxonomic units (OTUs) using CommunityM (https://github.com/dparks1134/CommunityM) under default settings.

### Metagenome assembly, binning, and bin dereplication

The raw reads generated from the Sunshine Coast Lake and Hikurangi Subduction Margin sediment DNA were first processed using SeqPrep (https://github.com/jstjohn/SeqPrep) under default settings to merge overlapping paired-end reads and trim adapters. Pre-processed paired-end reads were then assembled using metaSPAdes genome assembler v3.13.0 [63] with default settings. The raw reads from the Geothermal spring sediments were assembled using ALLPATHS [64]. Reads obtained from SRA were assembly using metaSPAdes with default settings. BamM (http://ecogenomics.github.io/BamM/) was then used to map sequences back to the assemblies. Next, binning was performed with uniteM (https://github.com/dparks1134/unitem) using selected methods (metabat_sensitive, metabat2, maxbin_107, maxbin_40 and groopM) under the default settings. CheckM [65] was then applied to calculate estimated completeness, contamination as well as strain heterogeneity. For metagenome-assembled genomes (MAGs) binned via multiple binning methods, the average nucleotide identity (ANI) was calculated, and MAG pairs with ANI greater than 99% were de-replicated by keeping the MAG with the highest quality, defined as completeness - 4 * contamination).

### Phylogenomics, rank normalisation and pangenomics

A total of 143 Asgardarchaeota genomes, including MAGs recovered from samples in this study, extracted from public SRA datasets, and downloaded from Genbank [66] with an estimated quality (completeness - 4 * contamination) over 40% were included in the downstream analysis. The multiple sequence alignment of selected MAGs was generated using gtdb-tk [67] based on 122 archaeal-specific marker proteins (**Table S2**). Maximum likelihood (ML) phylogenies for archaeal genomes were inferred using IQ-Tree 1.6.9 [68] under the LG+C10+F+G+PMSF model. Statistical support was estimated on a set of 1480 archaeal genomes (including 1377 non-Asgard archaea GTDB species representatives from GTDB release 05-RS95) using 100 bootstraps replicated under the same model (**Fig. 1,S4-5**). In addition, ML trees of trimmed alignments, from which we removed compositionally biased sites to increase tree accuracy for distantly-related sequences prior to concatenation, using BMGE [69]or Divvier [70], were evaluated with the same method (**Fig. S6-7**).

To further confirm the phylogenetic placement of Asgardarchaeota lineages, three additional ribosomal protein marker sets were used to create alignments: 16 ribosomal proteins defined in Hug et al [19], a subset of 23 proteins used by Rinke et al. [20] and a subset of 53 from the 56 top ranked archaeal marker proteins assessed in Dombrowski et al. [18]. Proteins were aligned to Pfam and TIGRfam HMMs using HMMER 3.1b2 (http://hmmer.org) with default parameters. The alignments were subjected to phylogenomic analysis using IQ-Tree 1.6.9 [68] under the LG+C10+F+G+PMSF model (**Fig. S8-10**). Bayesian trees were inferred with Phylobayes [71] for a subset of 44 genomes (incl. 34 Asgardarchaeota) under the CAT+GTR+G4 model (**Fig. S11**). Four independent Markov chains were run for ~43,000 generations. After a burn-in of 10%, convergence was achieved for all chains (maxdiff < 0.1). All phylogenetic trees inferred in this study are summarised in **Table S2**. Trees were viewed and annotated by iTOL [72].

The ranks of Asgardarchaeota lineages were normalised with PhyloRank (https://github.com/dparks1134/PhyloRank) based on the relative evolutionary divergence (RED) values, as implemented in the Genome Taxonomy Database (GTDB) [6, 7]; https://gtdb.ecogenomic.org/). Pangenomic analysis of selected Asgardarchaeota MAGs was conducted with Anvi’o version 6.2 [73] following its pangenomics workflow with option “–min-occurrence 2”.

To review the evolutionary relationship between Asgardarchaeota and eukaryotes, we used GraftM [74] for the identification of orthologues of 15 ribosomal proteins used in a previous studies [1, 22]. Eukaryotic hits were confirmed according to their NCBI annotation. The collected sequences for each marker gene were aligned with MAFFT v7.455 [75] and concatenated. The concatenated alignment was then trimmed by TrimAl v1.4 [76] with ‘-gappyout’ selection. Maximum-likelihood tree was calculated by IQ-TREE [68] under ‘LG+C60+F+G+PMSF’ model. Statistical branche support was calculated using 100 bootstraps under the same model.

### Proposed type material

MAGs proposal as type material were selected considering MIMAG standards [58] and following the recommended practice for proposing nomenclature type material [21].

### Metabolic annotation

Genes of all MAGs were predicted using Prokka [77] with the extensions “-kingdom archaea --metagenome” and annotated with EnrichM (https://github.com/geronimp/enrichM) against KEGG orthologs, EC, CAzy, Pfam and TIGRFAM databases for metabolic reconstruction. Predicted genes in major pathways were confirmed by querying the NCBI non-redundant (nr) protein database. Interpro IPR domains were assigned using InterProScan 5.31 [78].

#### Hydrogenase

We collected [NiFe]-, [FeFe]-, and [Fe]-hydrogenase sequences from the study of Greening et al. [79] to create a Blast database, which was used to query the 143 Asgardarchaeota genomes to search for potential hydrogenase genes. The sequence hits with e-values < 1e-20, scores >100, and sequence identities >30% were then submitted to HydDB [80] for further identification of hydrogenase subgroups.

#### Lipid membrane biosynthetic genes

KEGG orthologs of ester/ether lipid biosynthesis genes were used to investigate the potential of membrane lipid synthesis in Asgardarchaeota. To calculate the phylogenetic tree of glycerophosphoryl diester phosphodiesterase (UgpQ), we included genes used from a previous study of UgpQ phylogeny [27]. Eukaryotic UgpQ sequences were obtained from UniprotKB (http://www.uniprot.org) based on assignments to PF03009, including only sequences categorised as “Protein Existence [PE]" with the UniprotKB levels “Evidence at protein level” and/or “Evidence at transcript level”. Asgardarchaeota UgpQ homologues were identified with blastp [81] against KO K01126 by only retaining sequences with a maximum e-value of 1e-30. Collected UgpQ sequences were aligned using HMMER 3.1b2 (http://hmmer.org) against Pfam PF03009.

In addition, as the lipopolysaccharide ABC transporter genes were exclusively detected in Hodarchaeia MAGs, we inferred phylogenetic trees to rule out the possibility of misannotation. Sequences of lipopolysaccharide transport system ATP-binding protein (TagH, COG1134) and lipopolysaccharide transport system permease protein (TagG, COG1682) were collected from the NCBI conserved domain database Collected sequences together with Hodarchaeia hits for each COG were aligned using MAFFT v7.455 [75], respectively. Maximum-likelihood trees of UgpQ, TagH, and TagG were initially inferred by FastTreeMP [82] with Wag+Gamma model and subsequently with IQtree [68] under ‘LG+C60+F+G+PMSF’ model with 100 bootstraps.

#### ESP identification

High-quality Asgardarchaeota genomes (completeness>90%; <10% contamination; n=38) were selected to search for eukaryotic signature proteins (ESPs) listed in the annotation table in Zaremba-Niedzwiedzka et al. [1] The analysis was limited to high quality MAGs in order to minimize false negative hits. The resulting information was used to complete the ESP presence/absence table (**Table S4**). We used Prodigal [83] for gene prediction and hypothetical genes were annotated by InterProScan 5.31 [78] to screen for ESP homologs with certain IPR domains. As for ESPs denoted by the COG database, we downloaded sequences for each COG entry from the NCBI conserved domain database [84]. The COG sequences were passed to GraftM 0.13.1 [74] to create GraftM packages, which were then used to query Asgardarchaeota genes, with ‘graftM create’ and ‘graftM graft’ functions under default settings, respectively. Hits were further confirmed by blastp [81] against the NCBI non-redundant protein database (https://blast.ncbi.nlm.nih.gov/Blast.cgi).

#### Selenocysteine encoding system

We used Secmarker 0.4 [37] with the Infernal score threshold of 40 to detect the presence of tRNAsec in the Asgardarchaeota genomes and all archaeal and bacterial GTDB release 04-RS89 genus-dereplicated genomes. The detected tRNAsec sequences were aligned with MAFFT v7.455 [75] and trimmed by a minimum consensus of 40% [85]. Maximum-likelihood tree of tRNAsec was inferred using IQtree with 100 bootstraps under the VM+F+I+G4 model, which was selected by IQ-TREE’s ModelFinder module [68]. Seblastian [86] with default settings was applied to search for both selenocysteine insertion sequences and selenoproteins in Asgardarchaeota MAGs. The detected selenoproteins were verified by comparing the annotations to the corresponding Prokka-annotated genes with similar positions. Genes encoding enzymes responsible for selenocysteine biosynthesis and insertion were decided by annotation methods described above. Additionally, as the Thorarchaeota MAG “SMTZ1-83” is the only Asgardarchaeota genome proposed to encode SelA [36], we blasted the genes present on the contig (LRSK01000263.1) containing selA, using blastp [81] under NCBI non-redundant protein sequences database. The results are shown in **Table S12**, and reveal that this contig is most likely a contamination.

Since homologs of genes encoding SelB and SPS have been reported in archaeal, bacterial and eukaryotic genomes [35](Mariotti et al 2016), we hypothesised that these genes might be valuable to better understand the evolution of selenocysteine recoding. Bacterial and eukaryotic SelB and SPS sequences were selected and downloaded from UniprotKB (http://www.uniprot.org) to cover diverse taxonomic groups. Archaeal SelB and SPS sequences were collected from the order Methanococcales, two *Methanopyrus* genomes, and Asgardarchaeota, whose genomes were reported to be tRNAsec-positive. The collected gene sequences were aligned with MAFFT v7.455 [75] and trimmed by TrimAl v1.4 [76] with ‘-automated1’ selection. Maximum-likelihood trees were calculated by IQ-TREE [68] under ‘LG+C10+F+G+PMSF’ model with 100 bootstraps.

#### Pyrrolysine encoding system

The presence of tRNApyl in Asgardarchaeota MAGs was determined by Prokka 1.14.6 [77]. All genes of tRNApyl containing contigs of Thorarchaeia and Lokiarchaeota MAGs were compared against NCBI nr with blastp [81] to screen out possible contamination (**Table S16**).

Genes encoding enzymes responsible for pyrrolysine (Pyl) biosynthesis (PylS, PylB, PylC, PylD) and insertion (RF1) were detected by annotation methods described above. To explore the evolution of the Pyl system, we collected protein sequences of PylSBCD cluster genes (PylSc, PylSn, PylB, PylC, PylD) from the GTDB release 03-RS86 genus-dereplicated genomes. This was achieved by hmmsearch (Sean R. Eddy, http://hmmer.org) against HMM models of TIGR03912 (pyrrolysine--tRNA ligase, N-terminal region), TIGR02367 (pyrrolysine--tRNA ligase, C-terminal region), TIGR03910 (pyrrolysine biosynthesis radical SAM protein), TIGR03909 (pyrrolysine biosynthesis protein PylC), and TIGR03911 (pyrrolysine biosynthesis protein PylD). Homologs of PylB, PylC, and PylD that were not located on the same contigs were excluded, since these genes encode enzymes for pyrrolysine biosynthesis, and were only reported to be in close proximity. All genomes with at least two Pyl genes, which equals 50% of the required genes, were included in the downstream analysis. The collected sequences for each gene were aligned with MAFFT v7.455 [75] and trimmed by TrimAl v1.4 [76] with ‘-automated1’ selection. The PylS alignment was created by concatenating sequences of PylSn and PylSc. Maximum-likelihood trees were calculated by IQ-TREE [68] under ‘LG+C10+F+G+PMSF’ model with 100 bootstraps. Then we concatenated the above alignments in the order of pylSBCD, with the absence of certain genes represented by gaps. The contaminated alignment was trimmed by TrimAl v1.4 [76] with ‘-gt 0.4’ selection and further trimmed to exclude columns with less than 40% of consensus. Sequences with less than 80% remaining amino acids were removed, resulting in a final alignment of 62 protein sequences with 1103 columns. A maximum-likelihood tree was calculated with IQ-TREE [68] under ‘LG+C10+F+G+PMSF’ model with 100 bootstraps.

To search for Pyl-containing genes, we applied a strategy described previously [87]. In brief, we compared the annotation of each UAG-terminating CDS in all Hodarchaeia MAGs with the annotation of its downstream neighbouring CDS. In cases of matching annotations, both CDS were fused *in silico* as a unique CDS and predicted as potentially Pyl incorporating.

## Data availability

The raw reads and genome sequences from the metagenomes described in this study are available at NCBI under multiple BioProjects: PRJNA678545 (Sunshine Coast lakes) and PRJNA678552 (Hikurangi Subduction Margin).

## Acknowledgements

We would like to thank the shipboard scientists and crew of IODP Expedition 375 for collecting the Hikurangi Subduction Margin sediment samples, M. Chuvochina for etymological advice, and Brian K. for IT support. This work was funded by an Australian Research Council (ARC) Future Fellow Award (FT170100213) awarded to CR, and in part by the US National Science Foundation (DEB 1557042) and National Aeronautics and Space Administration (80NSSC17K0548).

## Notes

### Competing Interest Statement

The authors have declared no competing interest.

